# Layer-specific wide-field calcium imaging of neocortical activity

**DOI:** 10.64898/2026.04.28.721258

**Authors:** Dayra A. Lorenzo, Yasir Gallero-Salas, Matteo Panzeri, Anna-Sophia Wahl, Ariel Gilad, Christopher M. Lewis, Fritjof Helmchen

## Abstract

The mammalian neocortex is highly organized in local microcircuits and through long-range projection patterns between different regions. Since various features of local and long-range connectivity are determined by the cortical layer in which the respective neurons reside, understanding the flow of cortical information within and across layers is essential. Wide-field calcium imaging enables mesoscale functional mapping of genetically identified neurons across cortical areas. However, it has been applied primarily to superficial cortical layers, and systematic comparisons of wide-field signals across cortical layers are scarce. Here, we apply wide-field calcium imaging to different cortical layers using transgenic mouse lines with selective expression of GCaMP6f in layers 2/3, 5, and 6. We address several challenges of layer-specific wide-field imaging and provide possible solutions. First, to improve the registration of functional data to standard atlases, we demonstrate the benefit of layer-specific registration maps that are warped on the basis of the depth-dependent surface projections of the labeled cell populations. These maps help to assign the imaged calcium signals to the specific regions from which they originate. Second, we measure the depth-dependent blurring of wide-field fluorescence signals induced by light scattering and reveal stronger blurring in deep vs. superficial layers, in line with previous theoretical predictions from simulations. We used measured point spread functions to deconvolve single-whisker-evoked calcium signals in the barrel cortex and demonstrate improved signal confinement to individual barrel columns across layers. Finally, we investigate cross-regional functional connectivity during awake resting state periods for distinct layers. We find that mesoscopic functional connectivity is largely conserved between the cortical layers, with subtle differences for key regions of the default mode network (retrosplenial cortex and medial prefrontal cortex). Our approaches facilitate the comprehensive characterization of layer-specific cortico-cortical interactions, expanding wide-field calcium imaging as a powerful tool to investigate the layered organization of distributed brain dynamics.

## Introduction

The mammalian neocortex comprises multiple regions, each containing a local microcircuitry that is a variation of the canonical six-layer architecture (***Szentágothai, 1978; Douglas and Martin, 2004***). Long-range projections between the neocortical regions form a densely connected large-scale neural network (***Gămănuţ et al., 2018***), which enables perception, cognition, and adaptive behavior. A large diversity of neurons populates the cortical layers (***Ranson and Clark, 1959; Yao et al., 2023; Zhang et al., 2023; Chen et al., 2024***), forming intricate microcircuits that interconnect cortical and subcortical regions according to certain organizational principles (***Callaway, 1998; Douglas and Martin, 2004; Harris and Shepherd, 2015; D’Souza and Burkhalter, 2017; Vezoli et al., 2020***). In addition, different cortical layers make complementary functional contributions to the dynamics of local and distributed brain circuits (***Hubel and Wiesel, 1959, 1962; Krupa et al., 2004; Olsen et al., 2012; Chen et al., 2013; Li et al., 2015; Chen et al., 2015; Nandy et al., 2017; Gilad et al., 2018; Ayaz et al., 2019; Gallero-Salas et al., 2021; Esmaeili et al., 2021***).

The six-layered organization of the neocortex has been classically investigated in anatomical studies or by using penetrating electrodes (***Hubel and Wiesel, 1959, 1962; Mitzdorf, 1985; Mountcastle, 1997***). Modern optical methods, such as two-photon microscopy and wide-field fluorescence imaging, now in addition enable cell-type specific and depth-resolved physiological recordings (***Helmchen and Denk, 2005; Ayaz et al., 2019; Ren and Komiyama, 2021; Machado et al., 2022; Yamada et al., 2023; Murakami, 2024; Gilad, 2024***). Although wide-field microscopy does not provide optical sectioning, which is typically necessary for depth-resolved imaging, the use of transgenic mouse lines with layer-specific expression allows measurements of cellular activity in identified layers (***Madisen et al., 2015; Bethge et al., 2017; Musall et al., 2023; Mohan et al., 2023***). These advances provide exciting new opportunities to understand cortical dynamics on a large scale but at the same time in a cell-type-specific manner. Hence, there has been a surge in the application of wide-field imaging to study neocortical dynamics in mice (***Mohajerani et al., 2013; Wekselblatt et al., 2016; Allen et al., 2017; Makino et al., 2017; Gilad et al., 2018; Musall et al., 2019; Gallero-Salas et al., 2021; Nietz et al., 2022***). However, adoption and application of these approaches may still benefit from further characterization and refinement of the experimental methods.

To date, wide-field calcium imaging studies of cortical activity have primarily targeted superficial neurons in layer 2/3 (L2/3) due to their easy accessibility (***Gilad et al., 2018; Musall et al., 2019; Gallero-Salas et al., 2021; Esmaeili et al., 2021***). However, several recent studies have begun to use wide-field imaging to investigate specific neocortical populations not only in superficial but also deep layers, in particular layer 5 (L5), using genetic markers or projection-specific viral labeling (***Musall et al., 2023; Mohan et al., 2023; Heindorf and Keller, 2024; Pollak et al., 2025***). As the field moves toward exploring different regions and cortical layers, special challenges arise, particularly related to cortical depth. These concerns range from addressing basic anatomical differences across cortical layers, such as the curvature of the cortex, which is relevant to the parcellation of distinct areas, to evaluating and quantifying the issue of increased light scattering with cortical depth.

Here, we used transgenic mouse lines expressing the calcium indicator GCaMP6f in distinct neuronal populations of L2/3, L5, and layer 6 (L6) to perform wide-field calcium imaging of specific layers in awake mice. We introduce improved methods for layer-specific atlas registration and verify theoretical predictions of depth-dependent light scattering (***Waters (2020***)) by measuring layer-specific spatial kernels for deconvolution. We demonstrate the advantages of these improved methods for investigating across-layer differences in the spatial spread of cortical activation upon single-whisker stimulation and by analyzing layer-specific functional connectivity patterns between cortical regions (***Fox and Raichle, 2007; Ma et al., 2016; Mitra et al., 2018; Lake et al., 2020; Barson et al., 2020; Engel et al., 2021***). Our results suggest that the large-scale pattern of resting-state functional connectivity in pyramidal cells of the mouse dorsal cortex is largely consistent between layers. However, we do find subtle connectivity differences on a region-to-region basis, as well as differences in magnitude of these functional correlations across cortical layers.

## Results

### Anatomical considerations for top-view imaging

To measure hemisphere-wide calcium signals in different layers of the cortex, we performed wide-field calcium imaging of neuronal signals through the intact skull (***Silasi et al., 2016; Gilad et al., 2018***) using three distinct triple transgenic mouse lines that display specific expression of GCaMP6f in excitatory neurons of L2/3, L5, and L6, respectively (***Madisen et al., 2015***) (***Figure 1A-C***; Methods and Materials). To facilitate comparisons between individuals and statistical analysis, imaging data from individual mice are typically registered using a common anatomical reference space. The required spatial transformation can be done using linear or non-linear registration methods based on anatomical and functional landmarks. The Allen Mouse Common Coordinate Framework (Allen CCF) (***Oh et al., 2014; Harris et al., 2019; Wang et al., 2020***) has become the reference space of choice for wide-field imaging data, allowing comparison of results not only within individual studies but also between studies (***Allen et al., 2017; Musall et al., 2019; Esmaeili et al., 2021***). However, the reference map for the dorsal cortex is typically derived from the bird’s-eye view of the surface outlines of the cortical areas. Although this map is well suited for superficial L2/3, it does not account for the curvature of the cortex which distorts the top view projections for deep layers. To address this issue, the registration approach needs to be adjusted for deeper cortical layers. Functional landmarks, such as those derived from sensory-evoked activity maps, can be particularly beneficial in this process.

**Figure 1.**
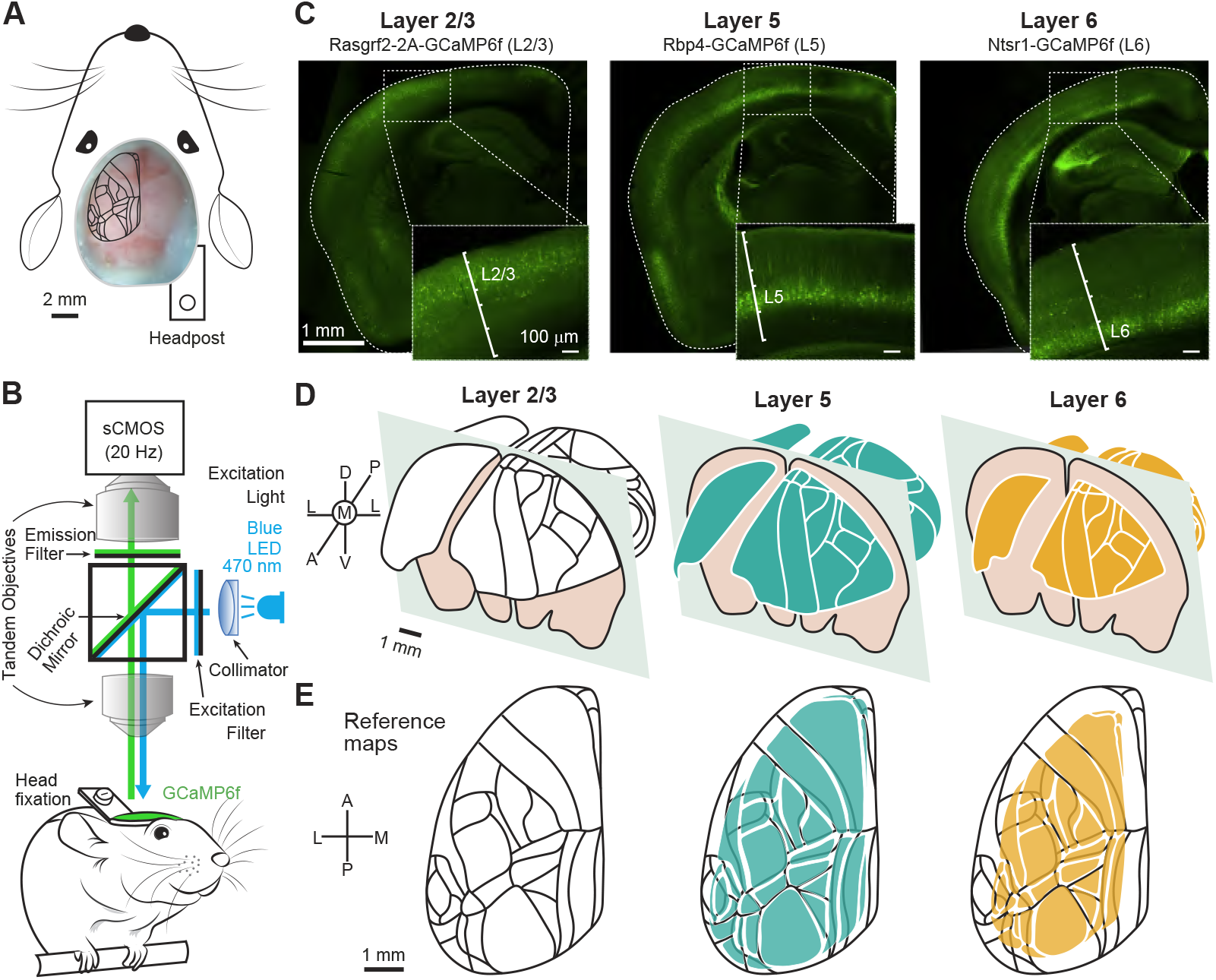
Anatomical considerations for registration of mesoscale imaging data. (**A**) Intact-skull wide-field preparation with registration pattern superimposed. (**B**) Schematic of wide-field imaging system. (**C**) Coronal slices of GCaMP6f-expressing triple-transgenic mouse lines specific to L2/3, L5, and L6. (**D**) Three-dimensional schematic of the layer-specific reference maps based on the Allen CCF for L2/3, L5, and L6. (**E**) Transversal top view of panel D superimposed on the L2/3 reference map (black lines).

In wide-field calcium imaging, the observed field of view is a two-dimensional projection, most commonly from above the dorsal cortex. From this dorsal perspective, neurons in distinct layers are not axially aligned but overlap to varying degrees due to the curvature of the brain surface (***Figure 1D***). Because of this curvature, the signals in the top projection view vary systematically depending on which layer they originate in, with deeper layers appearing more confined from the camera’s perspective. To take this into account, we calculated the horizontal plane projection for the different layers based on the Allen CCF (***Wang et al., 2020***). A comparison of the resulting reference maps shows that the map shrinks as the depth of the considered layer increases (***Figure 1E***). Thus, for registration of wide-field imaging data with layer-specific expression, we propose applying the appropriate layer-specific Allen CCF reference map.

### Layer-specific imaging of sensory-evoked activity maps

We applied layer-specific reference maps to align sensory-evoked functional maps in the three mouse lines. To drive specific sensory-evoked activities under light anesthesia, we presented the animal with a set of different sensory stimuli (***Figure 2A***; whisker, forelimb, hindlimb, visual and auditory stimulation) (***Mohajerani et al., 2013; Gallero-Salas et al., 2021***). In all mouse lines, we observed clear calcium signals (percent fluorescence changes, %Δ*F /F*) after stimulation and obtained activity maps from the Δ*F /F* values around the initial stimulus-evoked peak (mean of 1-s window after stimulus onset). For the different stimuli, calcium signals were largely confined to the respective primary sensory regions (***Figure 2B***). Slow negative signal components occurred with a delay, mainly after the stimulation period. In control experiments, we found that these negative signals represent hemodynamic responses, which did not, however, substantially affect our analysis of the initial response amplitude (***Figure 2–figure Supplement 1***). In particular, our data demonstrate that it is possible to record calcium signals from the L6 neuronal population through the intact skull using standard wide-field imaging, despite its substantial depth (***Figure 2B, bottom row***). Sensory-evoked functional maps allowed us to complement classic anatomical landmarks (that is, midline, bregma, and lambda) with functional landmarks and thus improve the registration of individual mouse data to our layer-specific Allen reference maps (***Figure 2B***; Materials and Methods). Registration with layer-specific maps was more accurate compared to using the standard L2/3 map and especially relevant for deeper layers. This is exemplified by registering images from example L5 and L6 mice with the L2/3 reference map, which led to obvious errors, particularly in lateral areas such as the primary auditory cortex (A1). These problems were avoided by using the appropriate L5 and L6 reference maps (***Figure 2C***).

**Figure 2.**
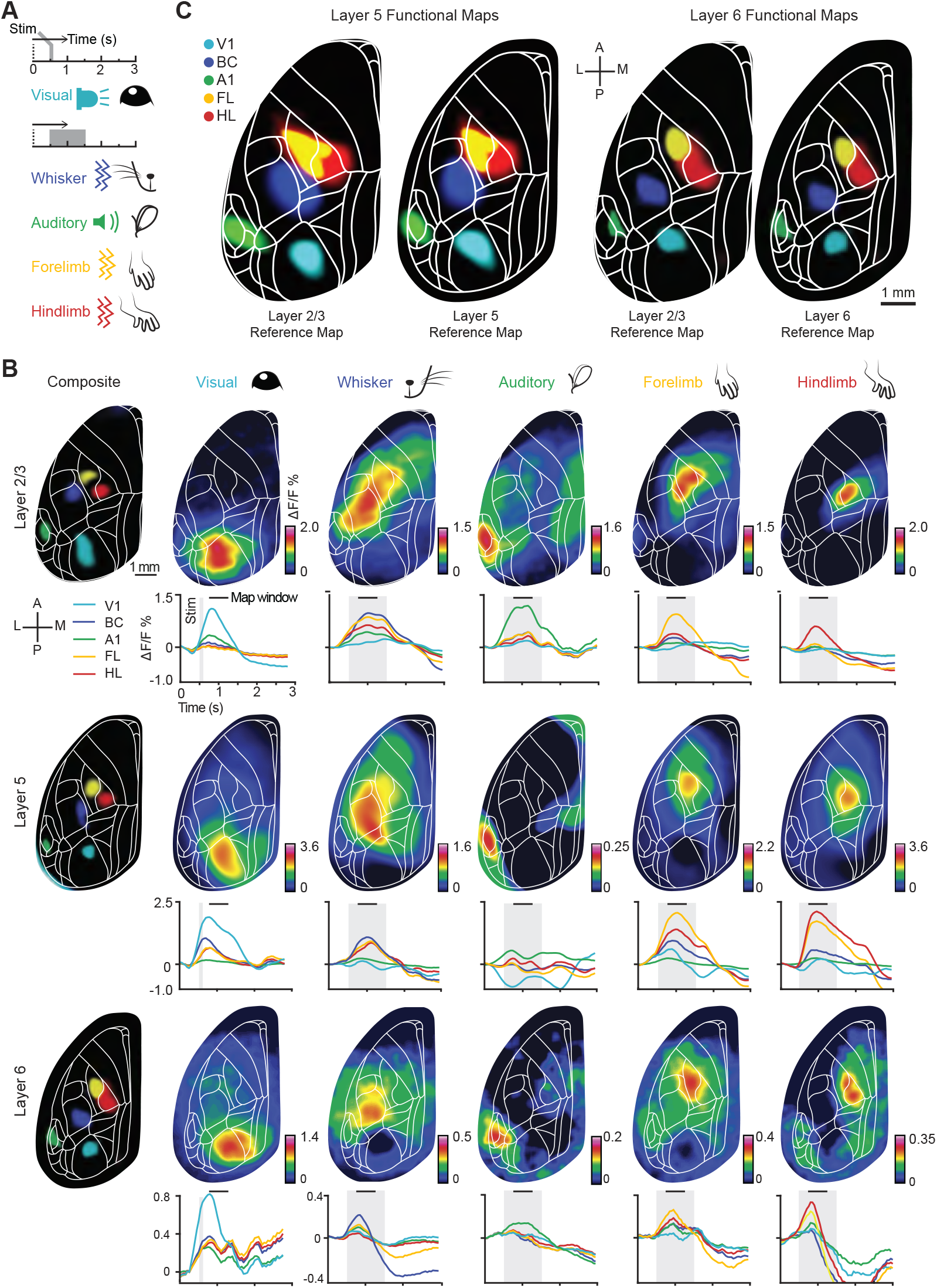
Layer-specific registration of sensory-evoked maps. (**A**) Schematic of wide-field imaging in anesthetized mice (left) and trial structure for the different sensory stimuli presented (right) (**B**) Average stimulus-evoked calcium activity maps (n= 10 trials), registered with layer-specific reference maps for L2/3, L5, and L6 (top to bottom row). Columns from left to right show the evoked activity maps for a 0.5-s window after stimulus onset for the five different sensory stimuli (min-max values of displayed fluorescence changes Δ*F /F* are indicated by color bars). Below each map the corresponding calcium traces are shown, with stimulation periods indicated as grey shade (100 ms for visual, and 1.0 s for the rest of the stimuli). (**C**) Benefit of registration using layer-specific reference maps exemplified for L5 and L6, respectively. Registering sensory-evoked calcium activity maps with the L2/3 reference map is suboptimal (e.g., for A1 and HL) but improves when the appropriate layer-specific reference maps are applied. **Figure 2—figure supplement 1.** Control functional mapping with hemodynamic correction.

### Depth-dependent effect of light scattering in mouse neocortex

Another consideration when imaging cell populations in different cortical layers is increased light scattering with depth (***Yizhar et al., 2011; Silasi et al., 2016; Yona et al., 2016; Waters, 2020***). Scattering of both excitation and emission light can affect the quality of imaging data, especially for deeper sources. In the case of wide-field imaging, scattering of excitation light is less relevant because fullfield illumination is used, and losses in signal brightness can be compensated for by increasing the illumination power. However, scattering of fluorescence emission light results in spatial blur, which reduces the lateral resolution of deep cortical sources when imaged from above (***Waters, 2020***). In principle, if the effects of scattering are known for different imaging depths, an appropriate point-spread function (or spatial kernel) can be used to deconvolve the images and thereby improve the resolution of deep sources.

To obtain depth-dependent spatial kernels for the deconvolution of wide-field imaging data, we measured the effect of light scattering by imaging a small artificial light source positioned at different depths inside the living mouse neocortex. We used an optical fiber with a prism at its tip, which we inserted horizontally from the lateral side into the cortex of anesthetized wild-type mice (n = 3 mice; multi-modal fiber with 70-*μ*m diameter, 50-*μ*m core diameter; targeted tip position from bregma: medio-lateral 1.5 mm, anterior-posterior -0.5 mm). We coupled 530-nm LED light into the optical fiber and ensured that the prism directed the light 90° upward through the cortex and the skull towards the imaging objective (***Figure 3A***). By sequentially inserting the fiber into the brain at different depths (from deep to superficial) while maintaining a constant light intensity, we could determine the attenuation and broadening of depth-dependent signals (***Figure 3B***). The peak intensity was reduced by 82 ± 3% (mean± SEM, n = 3) when imaging the fiber at a cortical depth of 1.0 mm, compared to a depth of 0.2 mm (***Figure 3C and D***). Normalizing the wide-field images of the quasi-point source to the maximum value obtained at different depths, we revealed increased spatial blurring with depth due to light scattering (***Figure 3E***). Because of signal loss and spatial blurring, deep images were particularly vulnerable to artifacts caused by vasculature and/or variations in the skull of individual mice (***Figure 3E***). We quantified the depth dependence of light scattering by calculating the radial profiles of the fiber tip images (***Figure 3F***). On average, the full width half maximum (FWHM, twice the half width of the radial profile) increased from 0.27 ± 0.03 mm at 0.2-mm depth to 1.01 ± 0.18 mm at 1.0-mm depth (mean ± SEM; n = 3; ***Figure 3G***). These results agree well with simulated depth-dependent scattering data (***Figure 3G***, see also ***Figure 3–figure Supplement 1***) obtained by applying previously published Monte Carlo simulations to model wide-field imaging in the mouse cortex (***Waters, 2020***).

**Figure 3.**
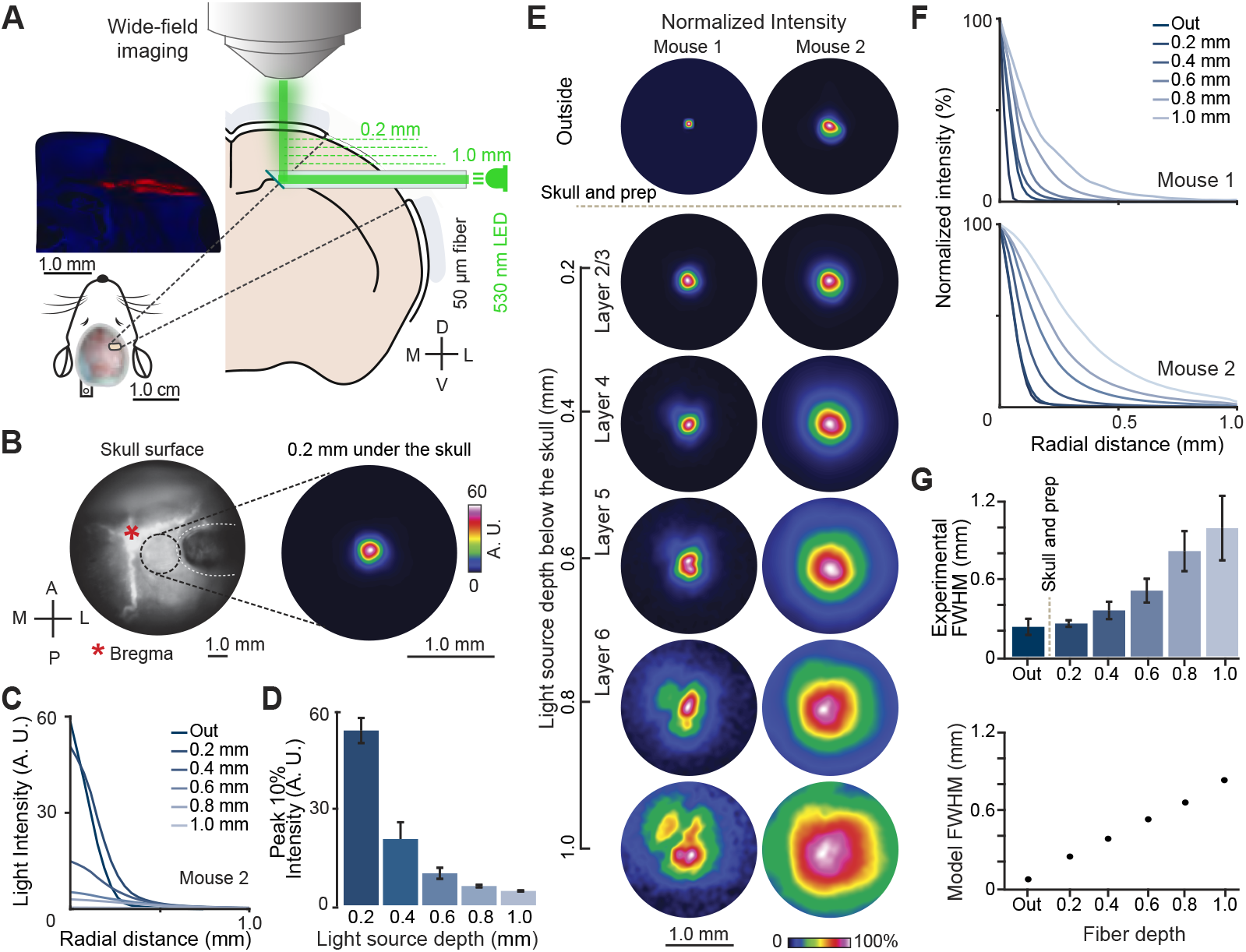
Wide-field imaging of light scattering across cortical depths. (**A**) Schematic of wide-field imaging of a point source of 530-nm light from an optical fiber with a 45° angle mirror at the tip, introduced from the side at different cortical depths in an anesthetized WT mouse. A histological coronal section showing the fiber insertion track (red dye) and DAPI stain is shown on top left. (**B**) Example bright field image of the skull surface (left) and fluorescence image of fiber-emission (right). Fiber tip at 0.2 mm depth. (**C**) Radial profiles of fluorescence intensity for an example mouse (outside and at 0.2-1.0 mm inserted depths). (**D**) Decrease in mean peak fluorescence intensity (peak 10% intensity) with depth. n = 3 mice; error bars are SEM. (**E**) Wide-field fluorescence images for two mice with the fiber source placed at different depths. Images are min-max normalized. (**F**) Radial profiles of the images shown in E, normalized to the central peak amplitude. (**G**) Top: FWHM of the experimentally determined radial profiles as a function of depth (n = 3 mice; error bars are SEM). Bottom: Corresponding FWHM values obtained from simulated data. **Figure 3—figure supplement 1.** Comparison of depth-dependent surface blurring due to scattering for experiment and simulations.

### Wide-field activity maps for single-whisker stimulation in L2/3 and L5 mice

We next aimed to demonstrate the benefit of the depth-dependent spatial kernels derived from our measurements of light scattering for deconvolution of functional cortical maps. We derived spatial kernels by fitting the average radial profiles of the quasi-point source at depths of 0.2-0.4 mm for L2/3 (n = 6) and 0.4-0.6 mm for L5 (n = 6) (***Figure 4A***). The well-known anatomy and physiology of the barrel cortex makes it a suitable model for studying calcium signals that originate from relatively small circumscribed regions in the cortex. For this experiment, we shaved all whiskers except the E1 and B1 whiskers and performed single-whisker stimulation for L2/3 and L5 mice under anesthesia (***Figure 4B***). We could detect single-whisker evoked calcium signals for both L2/3 and L5 neuronal populations, which we registered to our layer-specific reference maps (***Wang et al., 2020***) (***Figure 4C and D***). We then applied the Lucy-Richardson deconvolution algorithm (***Hanisch et al., 1996***) to estimate the true signal source using the depth-dependent spatial kernels derived from our scattering measurements. This approach improved the spatial acuity of the single-whisker-evoked activity maps compared to the original maps for both L2/3 and L5 (***Figure 4E and I***), evidenced by the narrowing of the radial profiles of the focally activated barrel columns in our images (***Figure 4F and G; J and K***) with significant decreases in FWHM for deconvolved images compared to original images for both L2/3 and L5 (***Figure 4H and L***). Thus, deconvolution with appropriate depth-dependent spatial kernels can improve the resolution of localized calcium signals.

**Figure 4.**
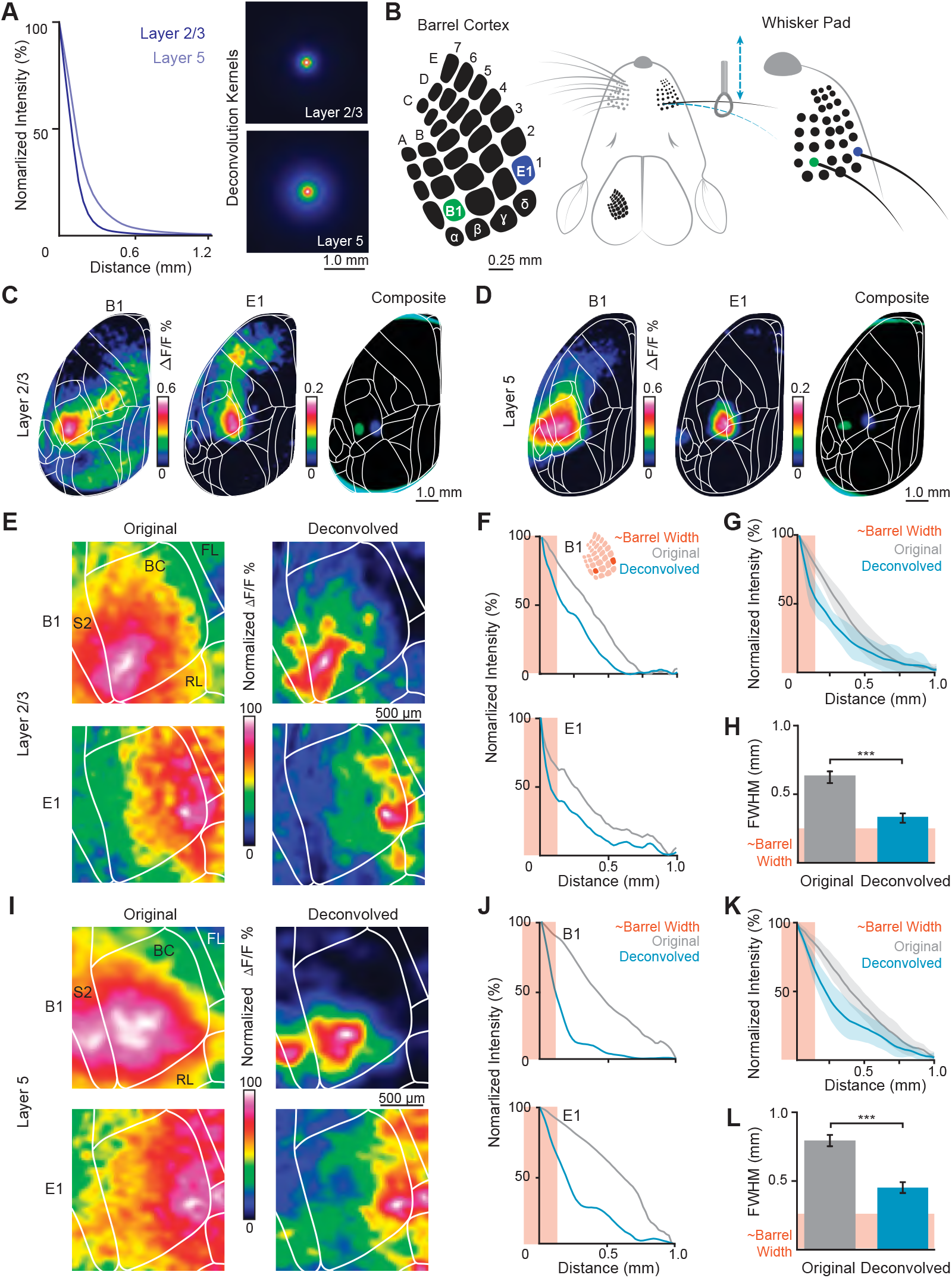
Deconvolution of single-whisker evoked maps using depth-dependent spatial kernels. (**A**) Radial profiles (left) and corresponding deconvolution kernels (right) for L2/3 and L5 generated from ***Figure 4*** (n = 3 mice per layer). (**B**) Schematic of single-whisker stimulation targeting B1 and E1 whiskers (right) and their corresponding barrel columns (left). (**C**) Example maps of B1 (left) and E1 (middle) stimulation in a L2/3 mouse (n= 20 trials); composite map (right) shows overlay of activation peaks. (**D**) Same as C for a L5 mouse. Zoomed-in examples of B1 (top) and E1 (bottom) stimulation in a L2/3 mouse: original images (left) and deconvolved images using the L2/3 kernel from A (right). Color scale: min–max normalized Δ*F /F*. (**F**) Radial profiles of original (grey; B1: FWHM = 0.72 mm, E1: 0.49 mm) vs. deconvolved (blue; B1: 0.34 mm, E1: 0.15 mm) images from E. Shaded box indicates approximate barrel width in L2/3. (**G**) Average radial profiles of original vs. deconvolved images (B1 & E1 pooled; n = 510 trials, 17 sessions, 2 L2/3 mice; error bars = SEM). (**H**) Mean FWHM values of profiles in G (original vs. deconvolved; error bars = SEM). (**I–J**) Same as E–F for a L5 mouse: original (grey; B1: 0.84 mm, E1: 0.87 mm) and deconvolved (blue; B1: 0.23 mm, E1: 0.30 mm) images, using the L5 kernel. (**K–L**) Same as G–H for L5 (B1 & E1 pooled; n = 360 trials, 2 mice). ^***^*p* < 0.001; Wilcoxon signed-rank test.

### Layer-specific assessment of resting-state functional connectivity

We next examined layer-specific functional connectivity of cortical regions during awake passive behavior, the so-called ‘resting state.’ Resting-state activity consists of dynamic patterns across the brain with inter-areal correlations that persist in the absence of explicit behavior or task participation. It has been widely examined in human neuroimaging and is believed to reflect previous experience, bias behavior, and correlate with disease state (***Engel et al., 2021; Fox and Raichle, 2007; Lewis et al., 2009; Baldassarre et al., 2012; Fox et al., 2009; Zhang and Raichle, 2010***). Wide-field calcium imaging has emerged as a promising tool for better relating resting-state networks observed with fMRI to the underlying neuronal and neuromodulatory activities (***Mohajerani et al., 2013; Vanni and Murphy, 2014; Matsui et al., 2016; Ma et al., 2016; Schwalm et al., 2017; Mitra et al., 2018; Bauer et al., 2018; Lake et al., 2020; MacDowell and Buschman, 2020; Lohani et al., 2022; Nietz et al., 2022; Collins et al., 2023; Benisty et al., 2024; Vafaii et al., 2024***). However, despite the increased use of wide-field imaging, few studies have investigated the layered structure of resting-state networks. Previous measurements of layer-to-layer dynamics during resting-state activity were performed with electrode arrays and therefore limited to specific regions of interest (***Mitra et al., 2018; Engel et al., 2021***). Here, our objective was to measure cortex-wide calcium dynamics in awake, resting mice with GCaMP6 expression specific to L2/3, L5, and L6, and to compare the resulting functional connectivity maps. Our results suggest that the overall structure of resting-state functional connectivity in pyramidal cells in the mouse neocortex generally shows a high degree of similarity across layers. However, we also point out subtle differences in connectivity patterns between layers.

We acquired calcium imaging data during the resting state in awake head-fixed L2/3, L5, and L6 mice (n = 6 mice per cohort, 200 to 300 trials per mouse, each lasting 10 s) (***Figure 5A***). We performed layer-specific registration and obtained GCaMP6 fluorescence traces for 26 cortical regions in the left brain hemisphere (***Figure 5B***) (Methods). We used a grouping scheme based on previously reported unidirectional anatomical connectivity, derived from targeted injections in layer-specific Cre lines and large-scale analysis of axonal projections (***Harris et al., 2019***). We monitored the face and body movements of the animals with videography. The region-to-region correlations of Δ*F /F* calcium signals (Spearmann rank correlation) showed high levels of correlation across all cortical regions for all layers (***Figure 5C***). In control experiments, we tested whether cross-regional correlations might have been affected by hemodynamic signals and found no significant confounding effect (***Figure 5–figure Supplement 1***).

**Figure 5.**
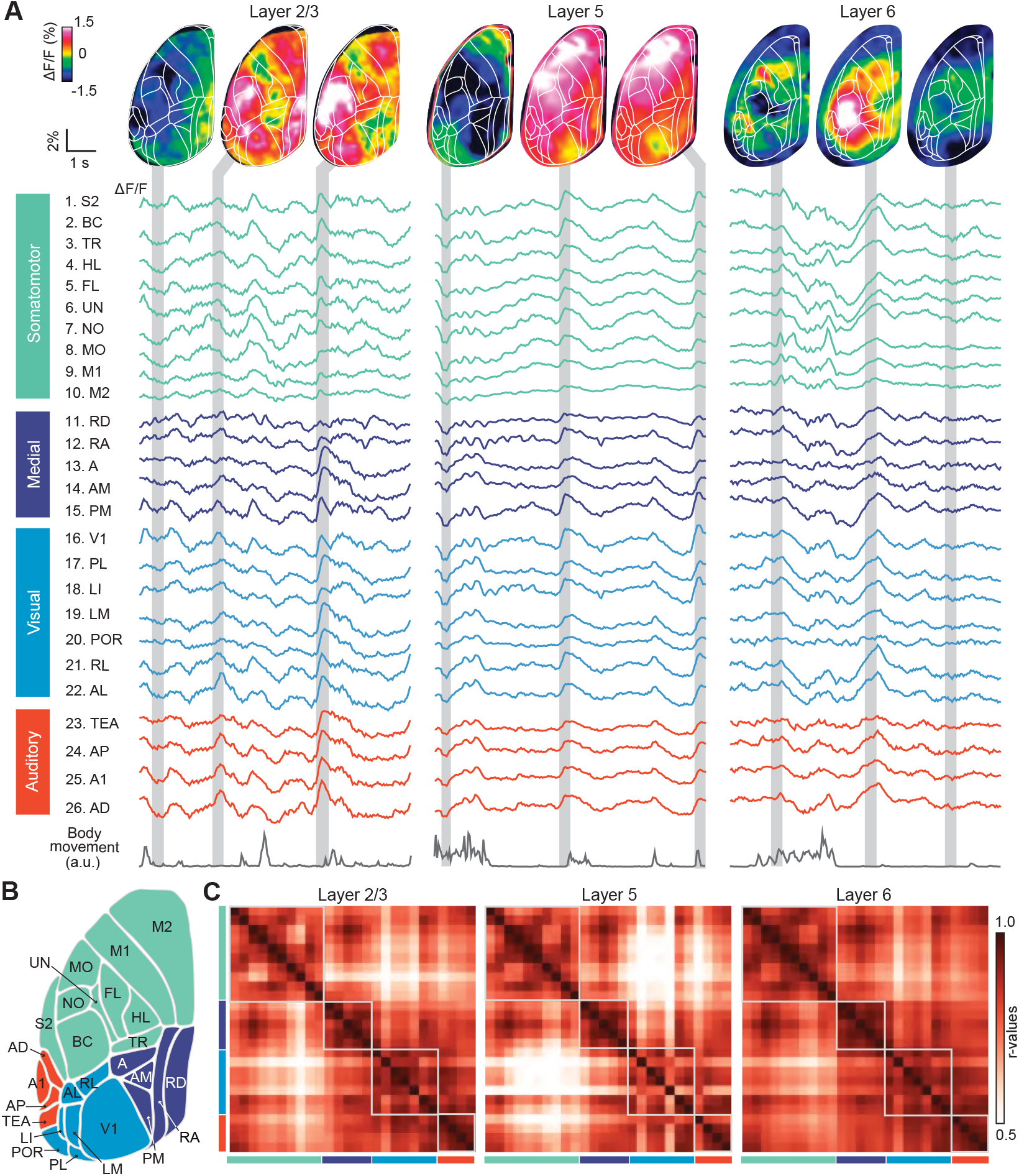
Layer-specific imaging of resting-state activity and cross-regional correlations. (**A**) Calcium activity maps (top) and raw Δ*F /F* traces in 26 cortical regions (below) from resting state measurements in a L2/3 (left), L5 (middle), and L6 (right) mouse. Example data from 10-s recording periods for each mouse. Gray-shaded areas indicate 0.3-s averaging periods, from which calcium activity maps were calculated. Bottom traces depict simultaneously recorded body movements. (**B**) Schematic of labeled regions in the left hemisphere with coloring of regional groups as in A. (**C**) Average region-to-region cross-correlation matrices for L2/3 (top; n = 6 mice), L5 (middle; n = 6), and L6 (bottom; n = 6). Regional grouping as in A and B; corresponding matrix modules are indicated by gray squares. **Figure 5—figure supplement 1.** Cross-regional correlations with hemodynamic control.

This high degree of correlation has been often observed in fMRI and large-scale cortical imaging, and it might be partially explained by global signals related to arousal and uncontrolled movements (***Schölvinck et al., 2010; Liu et al., 2017; Stringer et al., 2018; Musall et al., 2019; Aedo-Jury et al., 2020; Lohani et al., 2022; Collins et al., 2023***). To eliminate the impact of the global component of intrinsic brain activity on the correlations, we regressed the global mean signal from each region and performed a functional connectivity analysis on the residual time series. Global signal regression (GMR) is widely used in fMRI studies and has been shown to improve the identification of multiple resting state networks (***Fox et al., 2009; Murphy and Fox, 2017; Li et al., 2019; Satterthwaite et al., 2019; Lake et al., 2020***). After GMR, the functional connectivity patterns for all mouse lines showed more distinct regional segregation and a higher degree of anticorrelation between regions, especially between the regions of the somatomotor cortex and the primary sensory and association regions (***Fox et al., 2009; Murphy and Fox, 2017***) (***Figure 6A***). We verified in control mice that the connectivity matrices were not confounded by hemodynamic signals (***Figure 6–figure Supplement 1***). Across the three layers, functional connectivity patterns showed high similarity (***Figure 6A***) and were consistent with the anatomical connectivity patterns reported previously (***Harris et al., 2019***) (***Figure 6B***).

**Figure 6.**
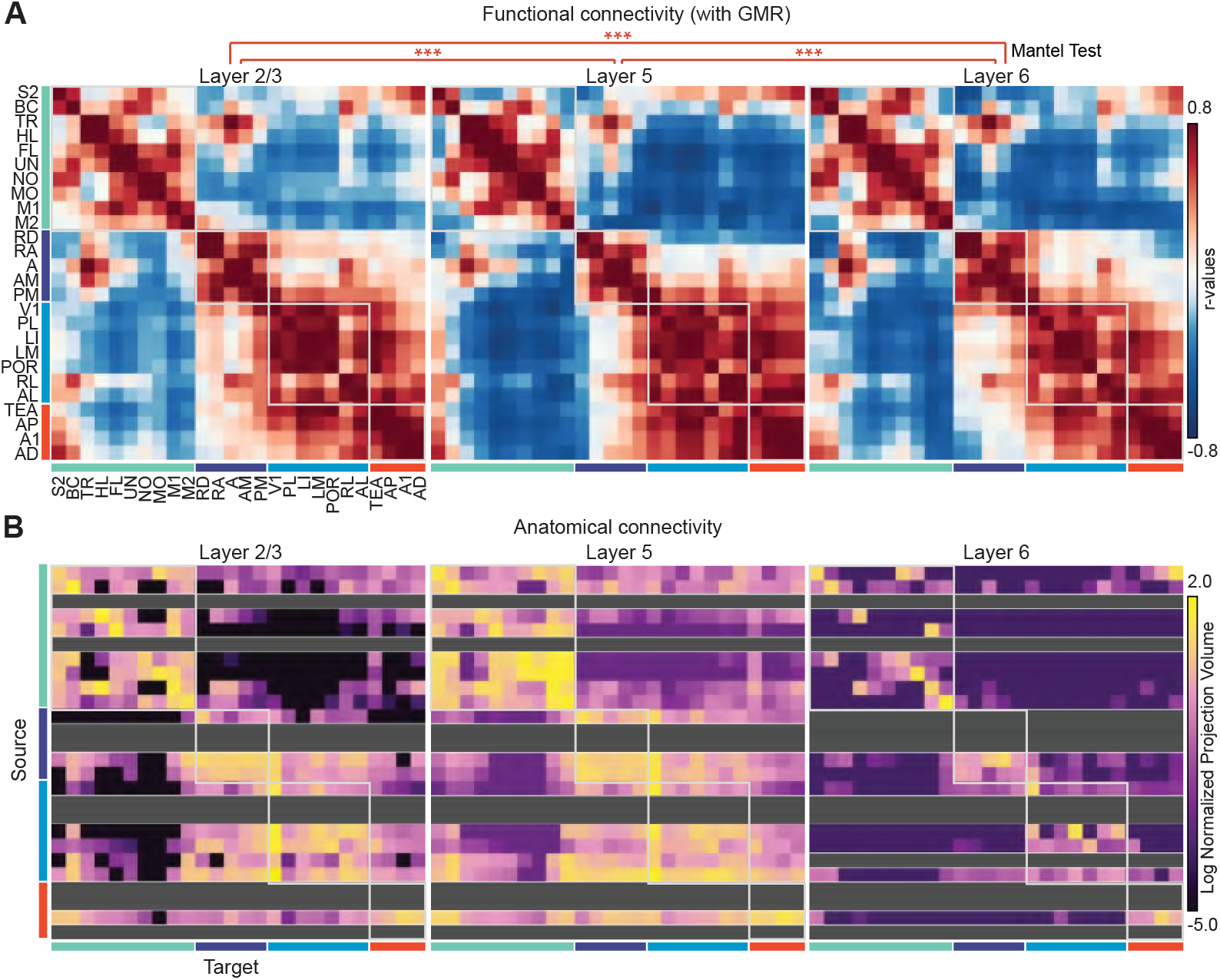
Layer-specific functional connectivity and anatomical comparison. (**A**) Average functional connectivity matrices for L2/3 (left; *n* = 6), L5 (middle; *n* = 6), and L6 (right; *n* = 6), showing cross-correlations of GMR-corrected Δ*F /F* traces across 26 cortical regions during rest. Matrices were highly similar (Mantel–Spearman: L2/3–L5, *r* = 0.96; L2/3–L6, *r* = 0.93; L5–L6, *r* = 0.93; 500 permutations; all ^***^*p* < 0.001). (**B**) Anatomical connectivity matrices for the same 26 regions in three mouse lines with layer-specific expression in the source regions. Each row contains the connection strength from the source region to all target regions (columns). Connection strength is expressed as log_10_-transformed normalized projection volumes (NPV). Axonal projections to all layers were considered in the target regions. Rows for which experimental data were missing (often because of low Cre expression) are indicated in grey. Data from (***Harris et al., 2019***). **Figure 6—figure supplement 1.** Functional connectivity with hemodynamic control.

To further assess potential differences in functional connectivity across cortical layers, we calculated pairwise contrasts between functional connectivity matrices from the different layers. This analysis revealed multiple region-to-region pairs with significant layer-dependent differences in correlation magnitude based on GMR-corrected traces (FDR-adjusted *p* < 0.001, 10,000 permutations; n = 10, 12, 10 for L2/3, L5, and L6, respectively; 6 mice per cohort). Interestingly, these differences were clustered topographically (***Figure 7A***), suggesting some organizational principles of layer-specific interregional interactions. For example, visual areas showed higher correlations with medial or frontal areas in L2/3 compared to deeper layers. However, within the somatomotor network, L2/3 showed lower functional connectivity. The total number of pairs of regions with significant differences varied between layers and within the cortex: It was more uniform between L5 and L6, and the posterior medial region RD showed the highest number of significant pairings (***Figure 7B***). The differences between L2/3 and the other layers were highest in the frontal premotor region M2 (***Figure 7B***). These observations align well with the prominent role that is assigned to both RD and M2 in the default mode network (DMN), a well-known resting-state network (***Raichle et al., 2001; Whitesell et al., 2021***). consistent with our findings, layer-specific differences in the anatomical connectivity profiles of these two areas were recently described (***Whitesell et al., 2021***).

**Figure 7.**
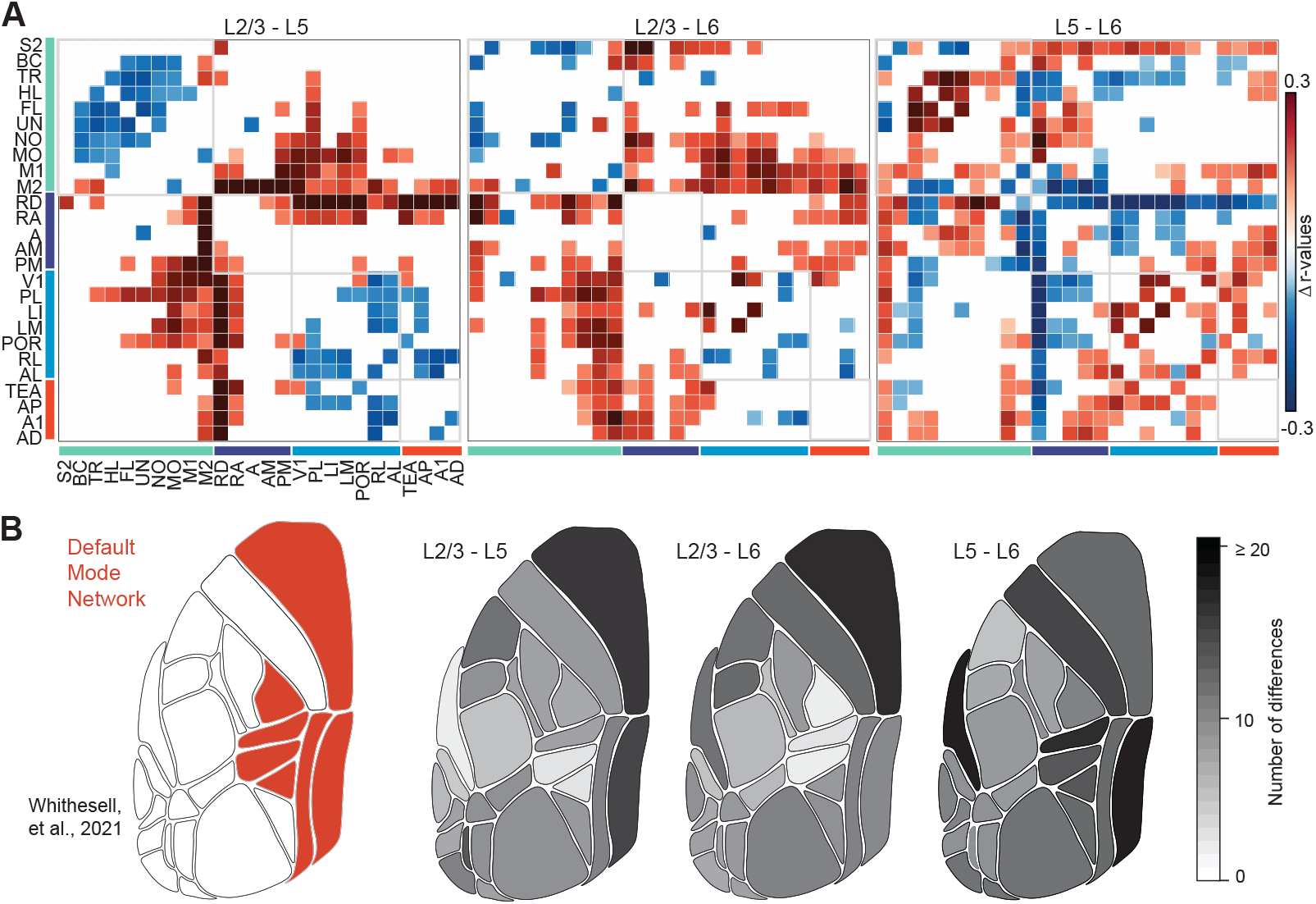
Layer differences in resting state functional connectivity. (**A**) Differences between the GMR-corrected functional connectivity matrices for L2/3 and L5 (left), L2/3 and L6 (middle), and L5 and L6 (right). Only areas with adjusted *p* < 0.001 are shown (FDR-corrected, 10,000 permutations). Regional grouping as in ***Figure 5***. (**B**) Left: Core regions of the default mode network as described in (***Whitesell et al., 2021***). Right: Topographical distribution of the number of region pairs with significant differences in functional connectivity (positive or negative) for each area comparing L2/3 to L5, L2/3 to L6, and L5 to L6 (from left to right).

## Discussion

We used wide-field calcium imaging to investigate layer-specific mesoscale activity across the dorsal cortex. By introducing layer-specific reference maps, we improved the accuracy of atlas registration, particularly for signals originating from deeper neuronal populations. In addition, we quantified the depth-dependent effects of fluorescence light scattering through the intact skull and demonstrated that the resulting spatial kernels can be used to enhance the spatial resolution of wide-field measurements. Finally, we assessed the layer-specific functional connectivity between brain regions. Although the overall structure of functional connectivity was largely conserved between layers during resting state, we also identified differences in connectivity between layers, most prominently in the retrosplenial cortex (RD) and secondary motor cortex (M2), regions associated with the default mode network. Our results demonstrate the great potential of applying wide-field calcium imaging in combination with specific labeling of neuronal subpopulations and appropriate analysis methods to characterize large-scale cortical dynamics.

### Considering the curved layering of the neocortex

The registration of wide-field calcium imaging data in a common atlas is important to compare data sets between individual subjects and between research groups. Although several suitable atlases are available (***Choi et al., 2024***), the Allen Mouse Brain Atlas is most widely used in the calcium imaging field. Most wide-field imaging studies have estimated regional boundaries by aligning their source data to the Allen CCF using affine transformation based on a few anatomical landmarks. This method is not ideal, because the precise position and identification of anatomical landmarks may vary between mice, with age, or with genotype (***Azimi et al., 2017***). In addition, the curved layering of the mouse neocortex is not taken into account, causing misalignment of the most lateral and medial regions, especially for deep sources (***Figure 2B***). To date, considering layer-specific registration of wide-field imaging has been largely overlooked, likely due to the relative scarcity of studies investigating specific layers or specifically labeled neuronal subtypes with cortex-wide imaging (***Esmaeili et al., 2021; Musall et al., 2023; Mohan et al., 2023; Heindorf and Keller, 2024; Pollak et al., 2025***).

Alternatively, some have proposed to define regions based solely on unsupervised clustering of functional data (***Barson et al., 2020; Xiao et al., 2021; Saxena et al., 2020***). Although such approaches are interesting and can allow the discovery of new functional units in an unbiased way, the resulting functional parcellation is likely to vary between individuals and, depending on the clustering criteria, may diverge from the histologically defined regions used to define classic anatomical targets (***Ren and Komiyama, 2021***).

Here, we demonstrate the importance of layer-specific registration, particularly for lateral regions when imaging deep layers (***Figure 2C***). We developed layer-specific reference maps for the Allen CCF (***Wang et al., 2020***) and based the alignment of data from individual imaging sessions with the reference maps on sensory-evoked functional maps, in addition to anatomical landmarks. This approach improves the identification of the regions and accounts for individual differences and brain curvature. It differs from the current wide-spread usage of only anatomical landmarks for linear fitting of the Allen CCF, and may improve the accuracy of registration and cross-study comparison of results even for L2/3. However, it should be noted that our registration scheme is based solely on the Allen CCF. Although this has become a common standard in the field, it displays notable differences from other popular atlases (***Gămănuţ et al., 2018***; **?**). Hence, layer-specific reference maps may also be created based on different atlases. Another factor that likely influences the registration of wide-field imaging data from deep sources is the degree to which cellular signals may originate from superficial compartments of deep-layer neurons, especially the dense apical arbors of L5 pyramidal neuron dendrites that extend into layer 1. Future work is needed to determine the extent to which more superficial dendritic signals can contribute to wide-field signals from L5 and L6 populations recorded at the surface, and how the mixing of superficial and deep sources affects the assignment of these signals to brain areas based on registration.

### Assessment of light scattering

Wide-field imaging is particularly sensitive to light scattering due to its lack of optical sectioning, especially when the emitted photons travel not only through the cortex but also through the skull (***Silasi et al., 2016; Gilad et al., 2018***). In vitro measurements and Monte Carlo simulations have indicated that light scattering can significantly reduce source localization, particularly for deep sources (***Yizhar et al., 2011; Waters, 2020***). Our measurements of the depth-dependent effect of light scattering in the brain of living mice complement and extend these results, and our extension of previous numerical simulations of scattering effects (***Waters, 2020***) may be particularly relevant for the field. We also demonstrate how derived deconvolution kernels can be applied to improve the spatial resolution of wide-field calcium imaging from deep cortical populations. These contributions might be especially relevant for small cortical regions, which could erroneously be reported to be active as a result of spatial signal spread from adjacent areas. Thus, deconvolution approaches should be particularly helpful in cases in which small regions are investigated or the topography of small populations within larger regions is of interest.

In our use case, deconvolution with empirical kernels increased the sharpness of wide-field calcium signals elicited from single-whisker stimulation. This approach could be used, for example, to study behavior-related whisker representations or map plasticity in the barrel cortex (***Minderer et al., 2012; Gilad, 2024***). Although our deconvolution kernels improved spatial resolution, they represent scattering properties estimated at the average depth of a given cortical layer; they do not fully account for the thickness of the layer nor for the projection of processes from neurons with a soma in one layer into other layers. It is also important to note that light scattering is not uniform, but can vary between anatomical structures (***Al-Juboori et al., 2013***) and between individual mice, depending on factors such as age and size. In the future, it will also be interesting to extend wide-field measurements to other wavelengths, for example, by using red-shifted indicators that exhibit reduced light scattering in tissue (***Dana et al., 2016; Bethge et al., 2017***).

### Layer-specific resting state connectivity

Finally, we demonstrate the utility of layer-specific imaging by comparing functional connectivity in the resting state as assessed for L2/3, L5, and L6 throughout the dorsal cortex. We found that the global structure of functional connectivity during the resting state is largely conserved between layers (***Figure 6***) but that the magnitude of region-to-region connection strength differs from layer to layer for a subset of regions (***Figure 7***), in particular for key components of the default mode network (***Figure 7B***). Our results agree with previous anatomical findings (***Harris et al., 2019***) and further confirm the close correspondence of functional connectivity in the resting-state with anatomical connectivity (***Honey et al., 2009; Grandjean et al., 2017***). Interestingly, the subtle differences in layer-specific connectivity that we observed in default mode network regions are in line with a recent study that combined fMRI and anatomical data in mice (***Whitesell et al., 2021***). This study reported differences in the pattern of layer-dependent connectivity, specifically between L2/3 and L5 pyramidal neurons in M2 and RD, the same areas that stood out in our study. Despite this converging evidence, it is important to note that the differences we observed were small and that the signal-to-noise ratio of surface-imaging data is expected to decrease with increasing depth of the imaged population. In addition, when imaging closer to the midline, the precise source of the signal becomes more uncertain because signals from various sources may overlap due to the strong curvature of the cortex at the midline. The largely conserved pattern of connectivity within all the studied layers may indicate that global connectivity is dominated by widely distributed components related to arousal, neuromodulation, and movement (***Musall et al., 2019; Lohani et al., 2022; Collins et al., 2023; Vafaii et al., 2024***). Within this stable connectivity pattern, subtle but meaningful variations emerge across cortical depth, possibly reflecting layer-specific contributions to large-scale networks. The precise functional relevance of these subtle differences is an interesting question for future research.

In summary, we have presented refined methods for layer-specific wide-field calcium imaging and demonstrated their utility in improving the localization of signal sources and in enhancing spatial resolution. Using these advances, we investigated the resting-state functional connectivity based on the correlation structure of spontaneous cortical activity in defined cortical regions. Distributed brain region networks are a central hallmark of large-scale brain organization, and activity maps are known not only to reflect underlying anatomical connectivity but also to relate to behavioral state, learned behaviors, cognitive demands, and alterations in disease states (***Liou et al., 2019; Mohan et al., 2023; West et al., 2024; Gilad, 2024; MacDowell et al., 2025; Nietz et al., 2025; Haley et al., 2026***). We anticipate the methods introduced here to facilitate a more precise investigation of layer-specific neural dynamics in the cortex and thereby advance our understanding of coordinated brain activity at the mesoscale.

## Methods and Materials

### Animals

All procedures were carried out according to the guidelines of the Swiss Federal Veterinary Office and were approved by the Zurich Cantonal Veterinary Office (licenses ZH211/2018 and ZH153/2019). A total of 26 adult male mice of 1-4 months age were included in this study. These mice were 6 wild-type (C57BL/6) mice and 20 triple-transgenic mice expressing GCaMP6f in excitatory neocortical neurons of layer 2/3 (n = 8; Rasgrf2-2A-dCre;CamK2a-tTA;TITL-G-CaMP6f), layer 5 (n = 6; Rbp4-Cre;CamK2a-tTA;TITL-G-CaMP6f), and layer 6 (n = 6; Ntsr1-Cre;CamK2a-tTA;TITL-G-CaMP6f). Triple-transgenic mice were generated by crossing double transgenic mice lines carrying the genes CamK2a-tTA and TITL-GCaMP6f with lines carrying the promoter genes Rasgrf2-2A-dCre, Rbp4-Cre and Ntsr1-Cre; for L2/3, L5 and L6, respectively (individual lines are available from The Jackson Laboratory as JAX *#* 016198, JAX *#* 024103, and JAX *#* 022864). Layer specificity is regulated through the expression of Cre and the layer-specific promoters mentioned above (***Bethge et al., 2017; Madisen et al., 2015***). Each of these lines contains a tet-off system, by which transgene expression can be suppressed upon doxycycline treatment. Here, we did not use this inducible system, so that neurons at the time of imaging presumably already expressed the indicator over several weeks, depending on the temporal profile of the activity of the promoters driving the expression of tTA and Cre. Note that the Rasgrf2-2A-dCre line contains an additional inducible system, given that the destabilized Cre (dCre) expressed under the control of the Rasgrf2-2A promoter needs to be stabilized by trimethoprim (TMP) to become fully functional. For TMP induction, TMP (Sigma T7883) was reconstituted in dimethylsulfoxide (DMSO, Sigma 34869) at a saturation level of 100 mg/ml and administered to mice by a single intraperitoneal injection 3 to 5 days before surgery (150 *μ*g of TMP/g body weight; 29 g needle; diluted in 0.9% saline solution). Imaging started earliest 10 days after TMP administration, by which the fluorescence has been reported to be stable (***Bethge et al., 2017***).

### Surgical procedures

For chronic wide-field neocortical calcium imaging, an intact skull preparation (without cover glass) (***Silasi et al., 2016***) was performed in all mice (***Figure 1A***). Surgeries were conducted under volatile anesthesia (2% isoflurane in pure oxygen) while maintaining body temperature at 37^◦^C. Full-body analgesia (0.2 mg/kg meloxicam; Metacam) was administered 20 min before surgery, and additional local analgesia (1% lidocaine) was applied to the skin before tissue removal. The head was shaved and cleared from hair (thioglycolic acid and potassium hydroxide cream; Veet), and the skin was disinfected (10% Povidone-iodine; Betadine) and removed above the dorsal skull. Dorsal muscles and connective tissue were carefully retracted laterally with a scalpel to expose the entire dorsal skull. Special care was taken to push the lateral muscles down the sides of the head to allow optical access to the auditory cortex without cutting the muscle (imaging field 6×8 mm^2^, spanning 3 mm anterior to bregma to 1 mm posterior to lambda, and from the midline to at least 5 mm laterally) (***Gallero-Salas et al., 2021***). After hemostasis, the skull was allowed to dry for 3–4 min, followed by application of an adhesive material (iBond; UV cured). A dental cement ‘wall’ (Charisma) was placed around the edges of the preparation, with the surrounding skin attached to the cement wall using tissue glue (VetGlue, 3M). Transparent dental cement (Tetric EvoFlow T1) was evenly applied across the imaging field. Finally, a metal post for head fixation was attached to the posterior skull with dental cement (***Figure 1A and B***) (***Silasi et al., 2016; Gilad et al., 2018; Gallero-Salas et al., 2021***). At the end of the experiment, mice were euthanized by a 5-min exposure to CO_2_. The brains were extracted, sectioned, and imaged using a confocal microscopy (FV1000, Olympus) to confirm fluorescent protein expression in the respective cortical layers (***Figure 1C***).

### Behavioral adaptation and experimental setup

Mice were handled daily for 5 days prior to surgery. Following recovery from intact skull preparation, earliest 5 days after surgery, we gradually adapted them to head immobilization in their home cage over a five-day period. Habituation began with 30-s immobilization periods and we progressively prolonged the duration to 15 min. After this habituation, the mice were introduced to the imaging setup and acclimated for 15 min on the day preceding the first recording session. At the end of the habituation period, the mice showed no resistance to the head-holding device. The awake mice were placed head-fixed under the wide-field imaging setup in darkness, with infrared LED illumination (940 nm, Monacor) from 30-cm distance. Wide-field calcium imaging was performed in 10-s periods (trials), collecting approximately 360 trials per session (1 hour total recording). Concurrently, mice were monitored at 30-Hz frame rate with an infrared camera (The Imaging Source; DMK 22BUC03; 720 × 480 pixels). To track general body movements in resting-state sessions, regions of interest (ROIs) were selected around the snout, the left ear, and both forelimbs, and movement vectors were computed as frame-to-frame correlations (1–corr(*f*_*t*_, *f*_*t*+1_)) for each ROI. The vectors were then averaged across ROIs to obtain a general body movement index.

Functional mapping, single-whisker stimulation, and some hemodynamic control experiments were performed in mice under volatile anesthesia (1-1.5% isoflurane in pure oxygen). Body temperature was monitored and maintained at 37^◦^C. For wide-field imaging of cortical activity maps evoked by single-whisker stimulation, we trimmed all whiskers except B1 and E1 in two L2/3 mice and two L5 mice. The stimulated whisker was placed in a wire loop coupled to a loudspeaker-driven vibrating bar (40 Hz). Each trial lasted 4.0 s (80 frames): 0.5 s baseline, 1.0 s stimulus, and 2.5 s post-stimulus. Intertrial interval was 5.0 s.

### Wide-field calcium imaging

We used a custom-made tandem-objective microscope setup (***Ratzlaff and Grinvald, 1991***) to record hemisphere-wide cortical activity in the L2/3, L5, and L6 transgenic GCaMP6f mice (***Figure 1A***) (***Gallero-Salas et al., 2021***). The excitation pathway consisted of a blue LED (470 nm; Thorlabs, M470L3), an excitation filter (480/40 nm BrightLine HC), a collimator (Thorlabs, ACL50832U-A), and dichroic mirror (510 nm; AHF, beamsplitter T510LPXRXT) housed in a filter cube (Thorlabs, DFM2) to direct the light onto the left hemisphere of the mouse. The illumination power at the preparation was <0.1 mW/mm^2^ (measured with the Thorlabs energy console; PM400K3). The emission pathway included two objectives (Navitar; top objective, D-5095, 50 mm f/0.95; bottom objective inverted, D-2595, 25 mm f/0.95), the dichroic mirror positioned between them, an emission filter (514/30 nm BrightLine HC), and a CMOS camera (Hamamatsu Orca Flash 4.0) mounted on top of the system. The imaging field of view had a diameter of 9.8 mm, enabling coverage of most of the dorsal cortex within a single hemisphere (here, the left hemisphere). Movies of fluorescence images were acquired in 10-s periods at a resolution of 512 × 512 pixels and a frame rate of 20 Hz. At the beginning of each imaging session, a reference image of the skull and the blood vessel pattern was obtained under illumination with a green fiber-coupled LED (503 nm; Thorlabs, M530L4) to provide anatomical landmarks.

### Functional mapping and atlas registration

We registered individual mouse brains from the different cohorts (L2/3, L5, and L6) to layer-specific reference maps calculated from horizontal plane projections for the different layers in the Allen CCF (***Wang et al., 2020***). To improve registration, we performed mapping of sensory-evoked population signals (***Mohajerani et al., 2013; Gilad et al., 2018; Gallero-Salas et al., 2021***) to obtain functional landmarks in addition to anatomical landmarks on the skull (i.e., bregma, lambda, and the midline). Mapping experiments were performed in darkness under mild anesthesia, with depth of anesthesia monitored by muscular tone (assessed resistance against gently pulling the limbs). Each mapping session consisted of five blocks, each comprising 30 trials, using stimulation of the hindlimb, whiskers, and forelimb, as well as auditory and visual stimulation, respectively (performed in that order). Stimuli were delivered on the side contralateral to the imaging site (except for the auditory stimulus) and designed to evoke neuronal activity in the corresponding sensory cortices. For somatosensory stimulations (hindlimb, whiskers, forelimb), a loudspeaker-coupled vibrating bar was used to oscillate the appendages (40 Hz, 1-s duration). Auditory stimulation consisted of white noise or pure tones (4 kHz and 8 kHz, 1-s duration). Visual stimulation was delivered by a blue LED placed in front of the eye (100-ms duration; approximately 0^◦^ elevation and azimuth ^◦^ in the visual field) (***Gilad et al., 2018; Gallero-Salas et al., 2021***). Each trial lasted 4.0 s (80 frames), consisting of 0.5 s baseline, 1.0 s stimulus, and 2.5 s post-stimulus period. The intertrial interval was 5.0 s.

Sensory mapping yielded five functional spots that were used, together with the anatomical landmarks, as anchoring points to register each individual brain to a reference map using a third degree polynomial transformation (***Gallero-Salas et al., 2021; Gilad and Helmchen, 2020***). Using the Allen CCF, we extracted for 26 regions the horizontal plane outlines (top-view) for L2/3, L5, and L6, from which we generated the three layer-specific reference maps for the left hemisphere. Based on the Allen connectivity atlas (***Oh et al., 2014; Harris et al., 2019***) and our previous work (***Gilad and Helmchen, 2020; Gallero-Salas et al., 2021***), we subdivided and sorted the regions into four groups (***Figure 5B***):

- **Somatomotor areas**: Secondary somatosensory cortex (S2), Barrel cortex (BC; primary somatosensory whisker), Somatosensory trunk (TR), Somatosensory hindlimb (HL), Somatosensory forelimb (FL), Somatosensory undetermined (UN; zona incerta), Somatosensory nose (NO), Somatosensory mouth (MO), Primary motor cortex (M1), and Secondary motor cortex (M2).
- **Medial areas**: Retrosplenial dorsal (RD), Retrosplenial agranular (RA), Anterior (A), Anterior medial (AM), and Posterior medial (PM).
- **Visual areas**: Primary visual cortex (V1), Posterior lateral (PL), Lateral intermediate (LI), Lateral medial (LM), Postrhinal (POR), Rostrolateral (RL), and Anterior lateral (AL).
- **Auditory areas**: Temporal association area (TEA), Auditory posterior (AP), Primary auditory (A1), and Auditory dorsal (AD).

For the comparison of functional connectivity maps with anatomical connectivity (***Figure 6B***), we downloaded data from the Allen Institute (https://connectivity.brain-map.org; ***Harris et al. (2019***)) and extracted the anatomical connectivity plots for the same L5 and L6 Cre lines as used here and a comparable L2/3 Cre lines (Cux2-IRES-Cre line for L2/3, *n* = 21 mice; Rbp4-Cre-KL100 line for L5, *n* = 27; and Ntsr1-Cre-GN220 line for L6, *n* = 21 mice). Local injections of a Cre-dependent anterograde tracer (EGFP) in all source regions allowed analysis of layer-specific anatomical projections. Connection strengths are expressed as log_10_-transformed normalized projection volumes (NPV) from single experiments with virus injections in one of 26 source areas (***Harris et al., 2019***). Note that axonal projection volumes in the target regions were analyzed in a layer-unspecific manner.

### Hemodynamic control experiments

In two L2/3 mice we assessed possible hemodynamic confounding effects in sensory mapping and functional connectivity experiments. The intact-skull imaging preparation was identical to the procedure described above, but using a transparent cyanoacrylate glue (Zap-a-Gap, Pacer Technologies) instead of dental cement in the center of the preparation. Wide-field imaging was performed at 20-Hz frame rate, alternating between a 470-nm excitation LED for calcium-dependent fluorescence (Thorlabs, M470L3) and a 405-nm LED as an isosbestic control (Thorlabs, M405L4) (***Couto et al., 2021***). This resulted in a frame rate of 10 Hz per channel. Data were acquired at 512×512 pixel resolution, spatially filtered with a 5×5 pixel Gaussian kernel and binned to 256×256 pixels for further analysis. Slowly varying temporal trends in the data were removed via a high-pass Butterworth filter (2^nd^ order, cutoff frequency 0.1 Hz), applied pixel by pixel. The fluorescence of the 405-nm channel, *F*_405_, was rescaled 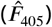 and subtracted from the 470-nm channel fluorescence (*F*_470_), yielding the hemodynamics-corrected fluorescence *F*_*corr*_. The scaling coefficients *β* were estimated for each pixel *i* by linear regression:

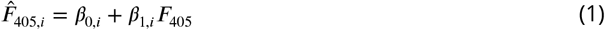

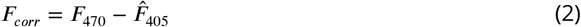

We then defined the uncorrected and corrected percentage fluorescence changes, (Δ*F /F*)_470_ and (Δ*F /F*)_*corr*_, respectively, for each pixel as follows:

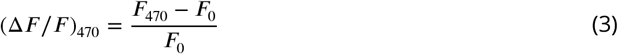

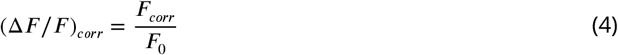

In the sensory mapping experiments, *F*_0_ was defined as the mean fluorescence of the 470-nm channel during the first 8 frames (0.8 s) of each trial. In the functional connectivity experiments, *F*_0_ was defined as the 10^th^ percentile of the 470-nm channel fluorescence. To ensure that all channels were comparable in scale, *F*_0_ was kept the same across channels. All additional preprocessing steps and subsequent data analysis, such as registration to the Allen atlas and calculation of functional connectivity matrices, were performed the same as for the main experiments.

### Measurement of light scattering

To experimentally assess the effect of light scattering in the mouse neocortex on wide-field imaging of different layers, we imaged a quasi-point source of light at different cortical depths and quantified the attenuation and spatial blurring of fluorescence signals in three anesthetized C57BL/6J mice (1% isoflurane anesthesia with body temperature maintained at 37^◦^C). Specifically, we used a multimode optical fiber with a 45^◦^ mirror tip as a point-like light source (50 *μ*m core diameter, 70 *μ*m total diameter, 0.22 NA, 4.0 mm length, SM3 receptacle; MFC 050/070 0.22 SM MA45, Doric). For stability, the fiber was mounted in a pulled glass pipette, leaving ∼0.5 mm of the tip exposed. The fiber was coupled to an LED light source (535 nm; Thorlabs, M530F2) via a multimode optical fiber cable (0.22 N.A.; M15L02, Thorlabs). Except for the very tip, the fiber was coated with the lipophilic dye DiI (Invitrogen; D282) to allow evaluation of the insertion site (***Figure 3A***). With the mirror at the tip pointing upward, the fiber was inserted horizontally from the side, targeting the hindlimb cortex (to avoid lateral curvature). We started with an insertion 1.0 mm below the cortical surface, recorded images, and then retracted and reinserted the fiber in 0.2 mm increments towards the surface of the cortex (depths of 1.0, 0.8, 0.6, 0.4, and 0.2 mm below the skull). Before insertion, the correct alignment of the fiber to direct light upward toward the microscope was verified by imaging the point source outside the skull and maximizing the intensity while rotating the fiber. A 2.0-mm craniotomy was prepared lateral to the position of the hindlimb cortex (0.5 mm AP, 4.0 mm ML from bregma) and the dura removed for fiber insertion. The fiber tip was advanced horizontally with a micromanipulator (LX20/M, Thorlabs). At each depth, we acquired 30 fluorescence images at 20 Hz: 10 baseline frames (light-off) and 20 frames with light on.

### Monte Carlo simulations

To simulate fluorescence light intensities at the brain surface we performed Monte Carlo simulations of photon scattering in neocortical tissue for emission sources at different depths. We adapted the Python Notebook provided in the previous study by Waters (***Waters, 2020***) and modified the model to include scattering through the skull and adapted the parameters for absorption and scattering to match our measurements with 530-nm LED light. Specifically, for 530-nm fluorescence photons in grey matter, we used an absorption coefficient of 0.5 mm^−1^, a scattering coefficient of 19 mm^−1^, and a value of 0.9 for the anisotropy factor in the Henyey-Greenstein phase function that determines the angular distribution of the scattered photons (***Jacques, 2013***). For the skull, we used a skull thickness of 0.25 mm, an absorption coefficient of 1.35 mm^−1^, a scattering coefficient of 26 mm^−1^, and a value of 0.9 for the anisotropy factor (***Soleimanzad et al., 2017***).

Surface distributions were simulated for photons emerging from a uniform distribution (50 *μ*m width) in the center of the field of view at various depths, mimicking the 50-*μ*m core of the optical fiber (NA 0.22) used for the experimental assessment of depth-dependent scattering. Photon trajectories were calculated using Henyey-Greenstein scattering. Light collection was modeled through a simulated 2x objective (NA 0.5) with a 22 mm field number, and the resulting spatial distribution of collected photons at the surface was characterized by fitting the exit coordinates to a Lorentzian distribution to determine the full width at half maximum (FWHM) of the light spread. The Python Notebook for the Monte Carlo simulations is available at …

### Quantification and statistical analysis

All analysis was performed using custom-written MATLAB scripts (MathWorks). Wide-field images (512 × 512 pixels) were downsampled to 256 × 256 pixels, and non-brain pixels were excluded. For functional mapping data, Δ*F /F* was calculated per pixel and per trial by normalizing the fluorescence changes to baseline *F*_0_ (average of 10 frames [0.5 s] preceding the stimulus). For resting state fluorescence data, baseline *F*_0_ was defined as the 15^th^ percentile of the fluorescence signal over the entire trial (10 min, i.e., 12,000 frames). (background subtraction?)

For light scattering measurements, reported images represent Δ*F*, defined as the average of light-on frames (20 frames) minus the average of light-off frames (10 frames), to correct for background fluorescence, such as skull autofluorescence. Radial profiles were calculated by identifying the pixel with peak intensity and averaging concentric rings, 1 pixel wide, around it, extending up to 53 pixels (2.014 mm). Direct point-source images were deconvolved using the Lucy–Richardson algorithm, with the optical fiber image obtained outside the skull serving as the kernel. Images were subsequently scaled using min–max normalization. Layer-specific deconvolution kernels were generated from average radial profiles of point-source images at the respective depths: 0.2–0.3 mm (L2/3; *n* = 3) and 0.4–0.6 mm (L5; *n* = 3). Radial profiles were revolved to generate point-spread functions, which were then applied to deconvolve L2/3 and L5 images obtained with single-whisker stimulation using the Richardson–Lucy algorithm.

For statistical analysis, we applied non-parametric tests throughout. Paired FWHM values (convolved vs. deconvolved) were compared using the Wilcoxon signed-rank test. Correlations were computed as Spearman rank coefficients. For matrix averages, Spearman *ρ* values were Fisher *z*-transformed and back-transformed for display. To compare connectivity matrices across layers, we performed Spearman–Mantel tests (500 permutations). Region-specific differences were evaluated using FDR-corrected permutation tests (10,000 permutations; *q* < 0.001).

## Acknowledgements

We thank Ladan Egolf and Philipp Bethge for managing triple transgenic mouse lines, Dubravka Göckeritz-Dujmovic for genotyping, Hansjörg Kasper for assistance with acquisition hardware and software, and Lazar Sumanovski for help with confocal imaging of histological samples. We thank Jack Waters for insightful discussions and help with the numerical simulations. This work was supported by grants to F.H. from the Swiss National Science Foundation (project grants 149858 and 170269) and the European Research Council (ERC Advanced Grant BRAINCOMPATH, project 670757), and to D.L. from the UZH Candoc Grant (Forschungskredit).

## Additional information

### Ethics

All procedures were approved by the Cantonal Veterinary Office in Zurich and carried out in accordance with guidelines of the Federal Veterinary Office of Switzerland.

### Resource Availability

#### Lead Contact

Further information and requests for resources and materials should be directed to the Lead Contact, Fritjof Helmchen.

#### Materials Availability

This study did not generate new unique reagents.

#### Data and Code Availability

Data and code supporting this study are available from Harvard Dataverse at: https://doi.org/10.7910/DVN/M7XIGH

## Author Contributions

D.A.L., C.L., A.G., and F.H. designed the experiments. D.A.L., C.L., Y.G.-S., M.P., and A.G. conducted the experiments. D.A.L and C.L. constructed the optical setup for wide-field imaging. D.A.L., C.L., Y.G.-S., and A.G. wrote the data analysis code. D.A.L., M.P., and C.L. analyzed the data. C.L.and F.H. performed numerical simulations. D.A.L., C.L. and F.H. wrote this article with input from the other authors.

---

**Figure 2—figure supplement 1.**
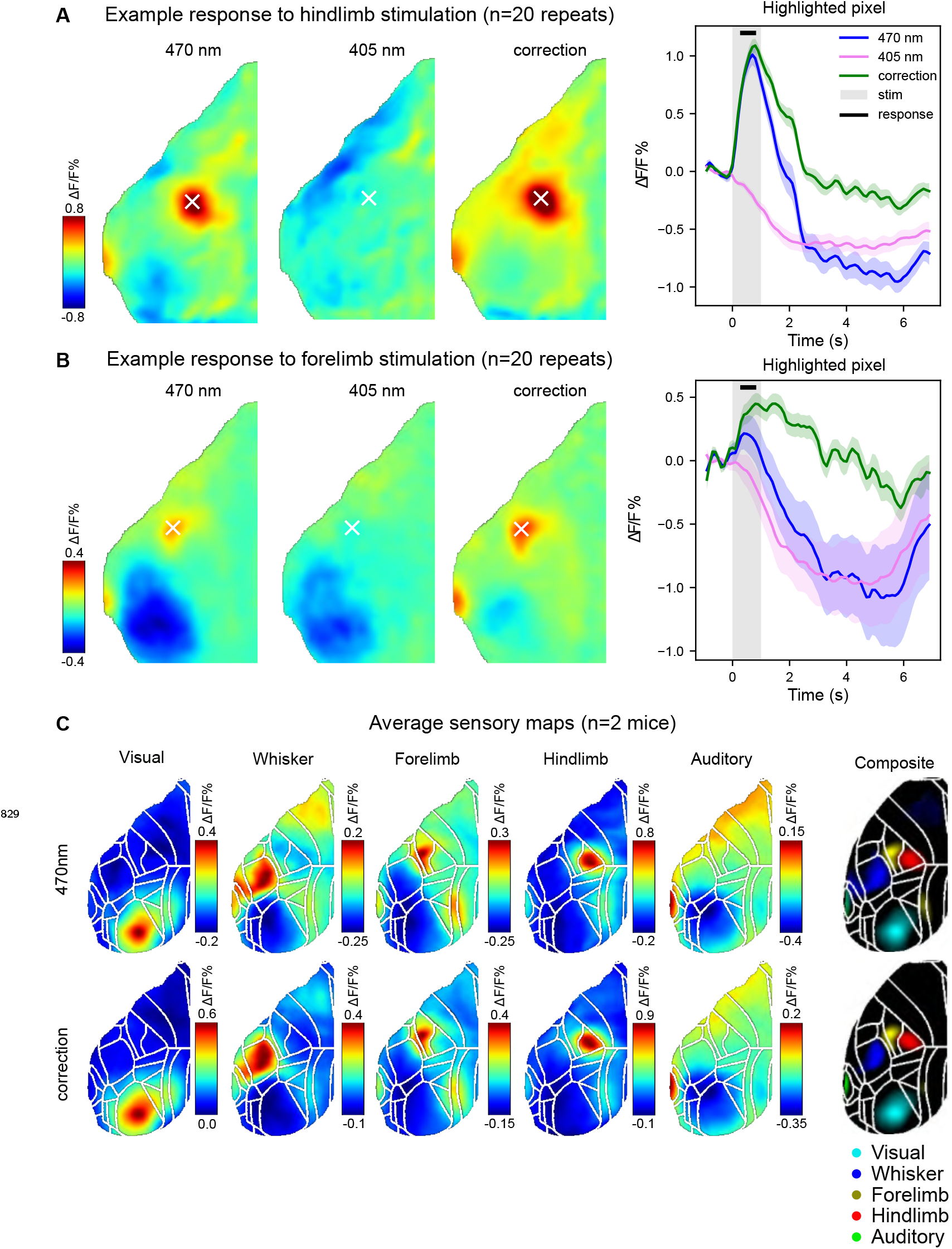
Control functional mapping with hemodynamic correction. (**A**) Example average functional maps upon hindlimb stimulation in a lightly anesthetized L2/3 mouse (n = 20 trials). Wide-field imaging was performed using dual-channel imaging with 2 excitation wavelength to correct for hemodynamic artifacts. Left: stimulus-averaged Δ*F /F* map with 470-nm excitation, maximizing GCaMP6f calcium sensitivity. Middle: stimulus-averaged Δ*F /F* map with 405-nm excitation, near the isosbestic (calcium-independent) wavelength of GCaMP6f. Right: corrected map Δ*F /F* obtained from the hemodynamic correction (Methods and Materials). Maps are calculated from the image frames within the stimulus window. (**B**) Same as (A) but with forelimb stimulation. (**C**) Comparison of Δ*F /F* maps for 470-nm excitation (upper row) and corrected Δ*F /F* maps (lower row) for all different types of stimulation (average from n = 2 L2/3 mice). Maps are aligned to the L2/3 reference map and the composite maps with all functional maps overlaid are shown on the right. No differences are apparent between 470-nm and corrected maps.

**Figure 3—figure supplement 1.**
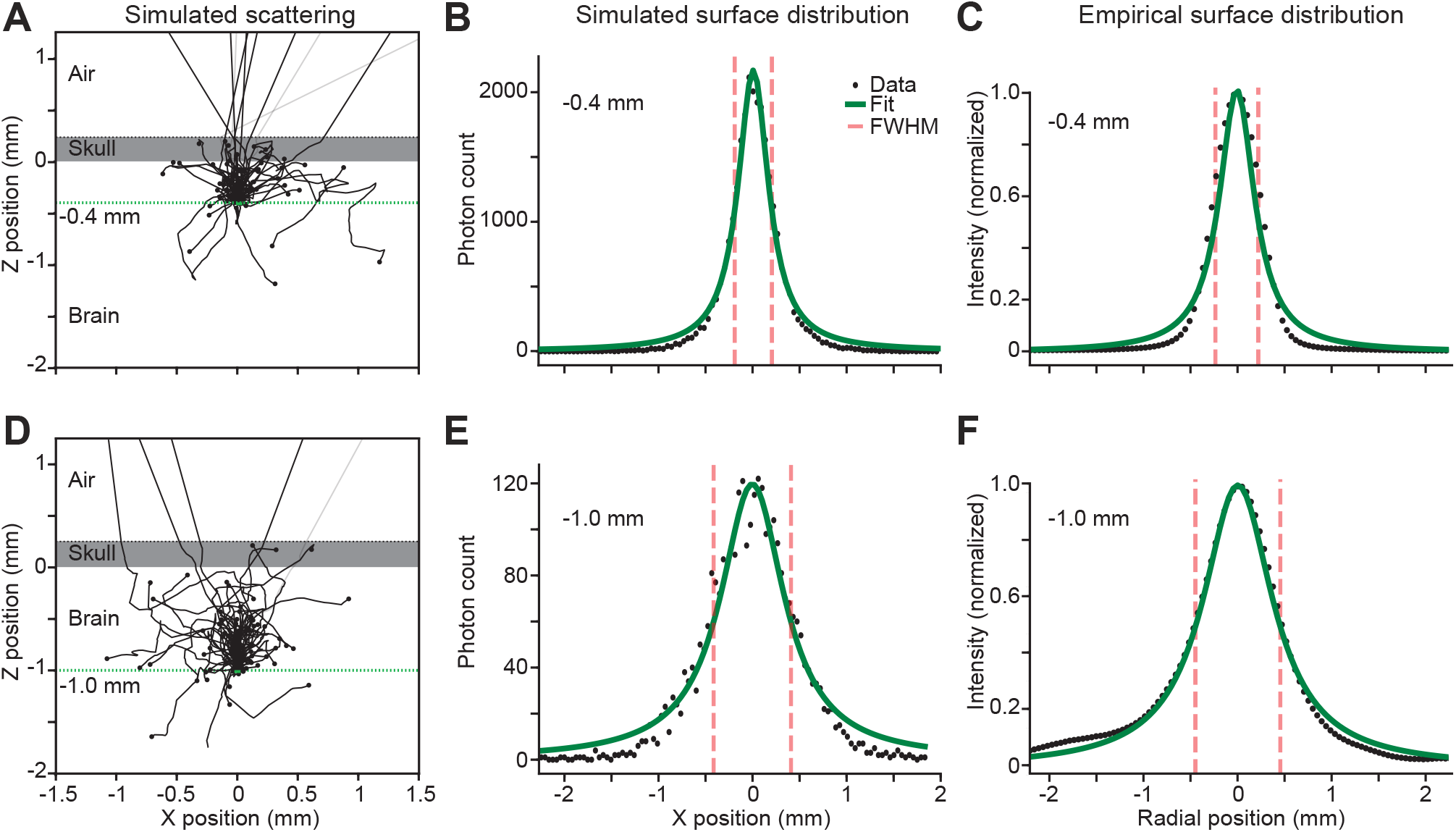
Comparison of depth-dependent surface blurring due to scattering for experiment and simulations. (**A**) Simulated scattering of 100 photons launched from a 50-*μ*m core diameter multi-modal fiber positioned 0.4 mm beneath the skull (dashed green line). (**B**) Distribution of photon detection across the surface based on simulations shown in (A) for 500,000 photons with a Lorentzian distribution fit (green line). (**C**) Distribution of experimentally measured light intensity across the surface of mouse 2 with a Lorentzian fit (green line). (**D**) Same as (A) but for 100 photons launched from a fiber 1.0 mm beneath the skull. (**E**) Same as (B) but for 1.0 mm fiber depth. (**F**) Same as (C) but for 1.0 mm fiber depth.

**Figure 5—figure supplement 1.**
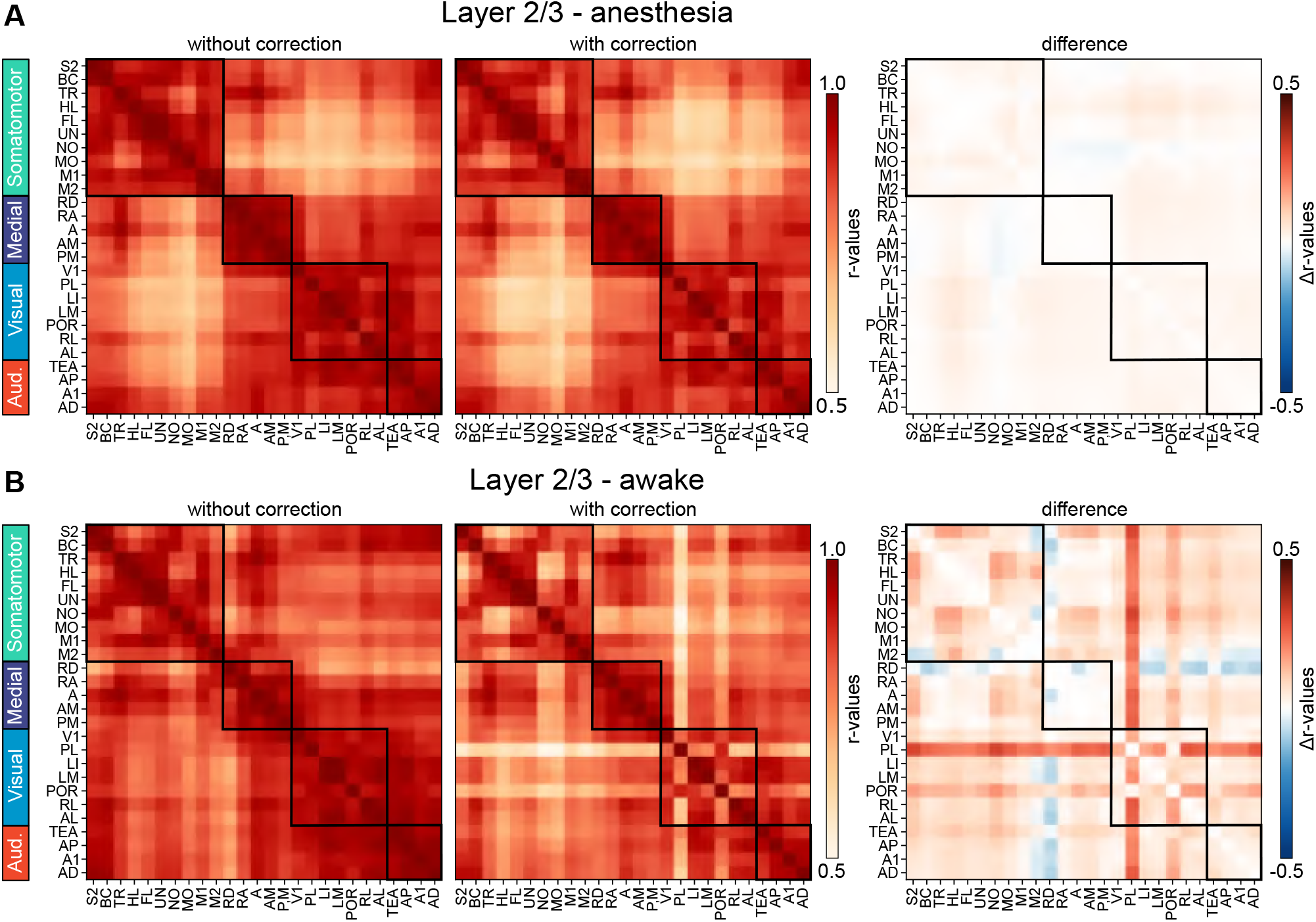
(**A**) Example cross-regional correlation matrices from an L2/3 mouse. (**A**) Forelimb stimulation under light anesthesia (n = 20 trials). Imaging was performed using dual-channel data (470 nm, 405 nm) to correct for hemodynamic artifacts. Left: correlation matrix from 470-nm channel without correction. Middle: correlation matrix after hemodynamic correction using the 405-nm channel. Right: difference matrix (uncorrected – corrected). Correlation matrices with and without correction were highly similar during anesthesia, as confirmed by Mantel–Spearman tests (*r* = 0.99, ^***^*p* < 0.001; 500 permutations). (**B**) Resting-state data from an awake L2/3 mouse. Left: correlation matrix from 470-nm channel without correction. Middle: correlation matrix after hemodynamic correction. Right: difference matrix. Correlation matrices with and without correction were also highly similar during the awake resting state (*r* = 0.77, ^***^*p* < 0.001; 500 permutations).

**Figure 6—figure supplement 1.**
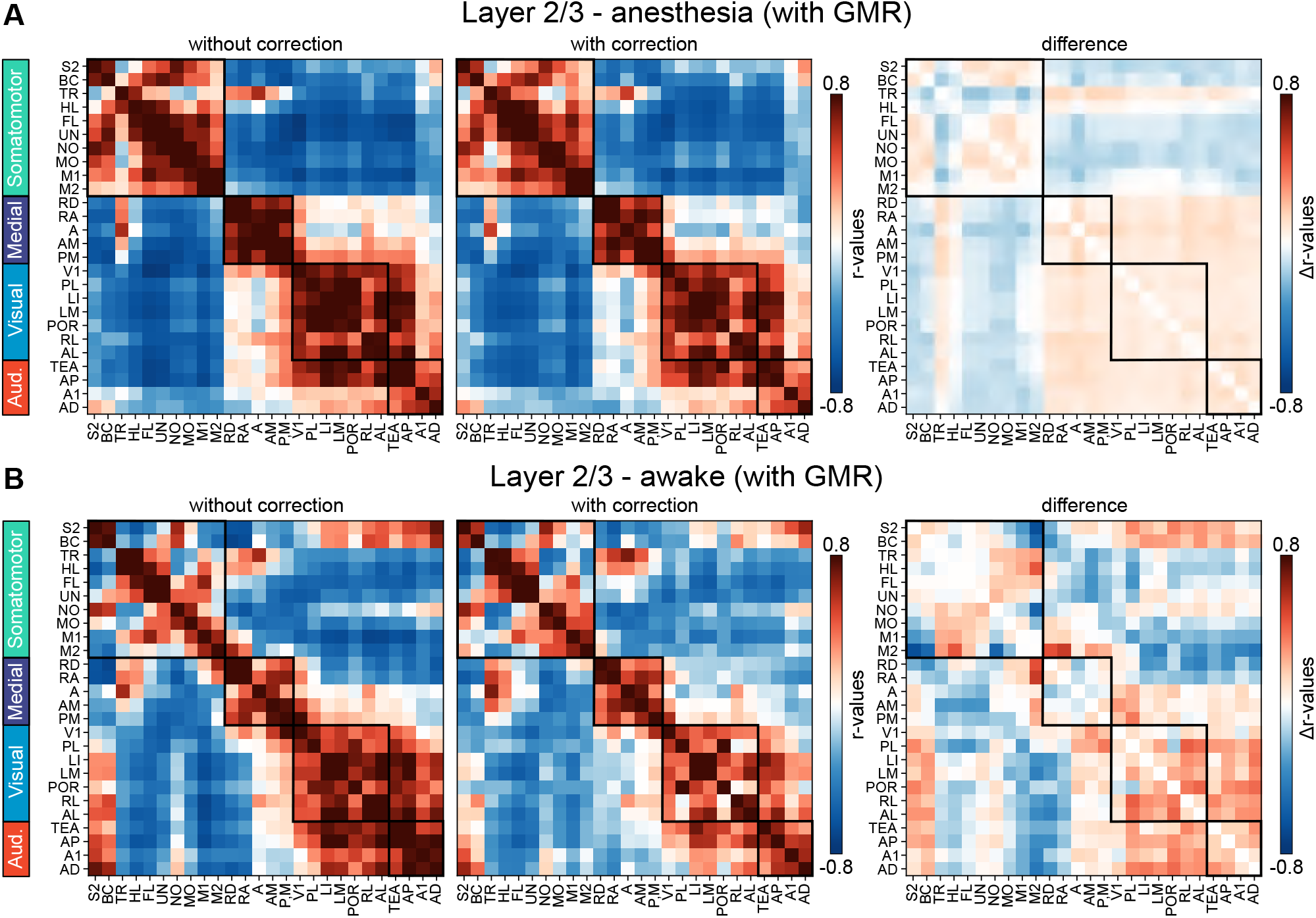
Functional connectivity matrices obtained with GMR using dual-channel imaging data (470 nm, 405 nm) from the same L2/3 mouse as in (***Figure 5, figure Supplement 1***). (**A**) Average functional connectivity matrices for forelimb stimulation under light anesthesia (n = 20 trials) without (left) and with (middle) hemodynamic correction using the 405-nm channel. Right: difference matrix obtained by subtracting the corrected from the uncorrected matrix. Connectivity matrices with and without hemodynamic correction were highly similar during anesthesia, as confirmed by Mantel–Spearman tests (*r* = 0.99, ^***^*p* < 0.001; 500 permutations). (**B**) Functional connectivity matrices from resting-state periods without (left) and with (middle) hemodynamic correction using the 405-nm channel. Right: difference matrix obtained by subtraction. Connectivity matrices with and without hemodynamic correction were also highly similar during awake resting state, as confirmed by Mantel–Spearman tests (*r* = 0.90, ^***^*p* < 0.001; 500 permutations).

